# Analysis of *Vibrio cholerae* genomes using a novel bioinformatic tool identifies new, active Type VI Secretion System gene clusters

**DOI:** 10.1101/526723

**Authors:** Cristian V. Crisan, Aroon T. Chande, Kenneth Williams, Vishnu Raghuram, Lavanya Rishishwar, Gabi Steinbach, Peter Yunker, I. King Jordan, Brian K. Hammer

## Abstract

**Background:** Like many bacteria, *Vibrio cholerae*, which causes fatal cholera, deploys a harpoon-like Type VI Secretion System (T6SS) to compete against other microbes in environmental and host settings. The T6SS punctures adjacent cells and delivers toxic effector proteins that are harmless to bacteria carrying cognate immunity factors. Only four effector/immunity pairs encoded on one large and three auxiliary gene clusters have been characterized from largely clonal, patient-derived strains of *V. cholerae*.

**Results:** We sequenced two dozen *V. cholerae* strain genomes from diverse sources and developed a novel and adaptable bioinformatic tool based on Hidden Markov Models. We identified two new T6SS auxiliary gene clusters; one, Aux 5, is described here. Four Aux 5 loci are present in the host strain, each with an atypical effector/immunity gene organization. Structural prediction of the putative effector indicated it is a lipase, which we name TleV1 (**T**ype VI **l**ipase **e**ffector **V**ibrio, TleV1). Ectopic TleV1 expression induced toxicity in *E. coli*, which was rescued by co-expression of the TleV1 immunity factor. A clinical *V. cholerae* reference strain expressing the Aux 5 cluster used TleV1 to lyse its parental strain upon contact via its T6SS but was unable to kill parental cells expressing TleV1’s immunity factor.

**Conclusion:** We developed a novel bioinformatic method and identified new T6SS gene clusters in *V. cholerae*. We also showed the TleV1 toxin is delivered in a T6SS-manner by *V. cholerae* and can lyse other bacterial cells. Our web-based tool may be modified to identify additional novel T6SS genomic loci in diverse bacterial species.

## Background

*Vibrio cholerae* is a globally dispersed, Gram-negative bacterium that naturally resides on chitinous surfaces in marine habitats. When ingested, some strains of *V. cholerae* can cause the fatal cholera diarrheal disease in humans. While relatively rare in developed countries, it is estimated that nearly 3,000,000 cases and 100,000 deaths from cholera occur annually, with the disease endemic to areas of Middle East and Southern Asia (1,2). Patient-derived strains (referred to as clinical strains) of *V. cholerae* possess virulence factors that help colonize the intestine and infect the human host (3). *V. cholerae* strains also possess other mechanisms to colonize hosts and persist in aquatic niches (4). An important defense employed by *V. cholerae* against other prokaryotic and eukaryotic cells is the Type VI secretion system (T6SS), a protein delivery system that punctures membranes of neighboring cells and delivers toxic effectors (Fig. 1a) (5,6).

**Figure 1.**
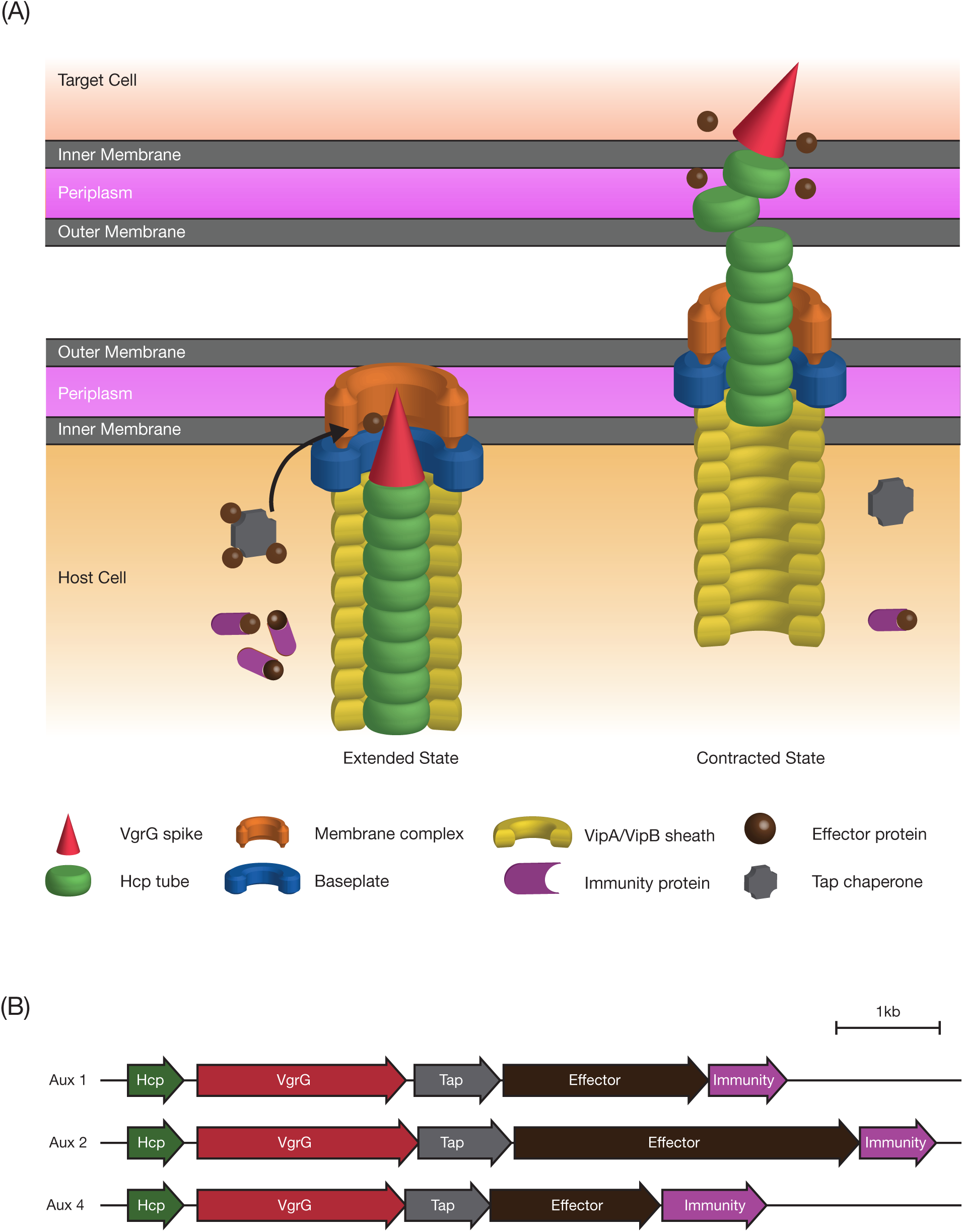
Type VI secretion system of *Vibrio cholerae*. A) Diagrammatic depiction of the T6SS apparatus assembly, contraction and disassembly in *V. cholerae.* The apparatus is composed of a membrane-spanning and a baseplate complex, an outer contractile sheath (VipA/B) and a needle complex (Hcp and VgrG). Effectors can interact directly with VgrG or PAAR proteins, may require chaperones for delivery on the apparatus or be carried as cargo in the T6SS apparatus. B) Aux clusters 1, 2 and 4 share a canonical *hcp, vgrG, tap, effector, immunity* gene organization in all strains where they are found.

Found in approximately 25% of all Gram-negative bacterial species, the T6SS apparatus consists of a membrane complex that spans both membranes and the periplasm of the host cell (7,8). A baseplate complex with homology to phage components attaches to the inner membrane and is thought to interact with other components of the apparatus (8,9). The T6SS functions through an ATP-dependent contractile mechanism facilitated by VipA/B sheath proteins (10–12). Hcp (Hemolysin-coregulated protein) hexamers form the inner tube of the apparatus and are exported into the extracellular milieu following contraction of the outer sheath (5,6,13,14). The tip of the apparatus is comprised of secreted VgrG proteins that interact with T6SS toxic proteins (called effectors) to aid in their delivery (15). PAAR proteins, found in some bacterial species harboring the T6SS, associate with VgrGs and are thought to sharpen the tip while also diversifying the cargo delivered by the T6SS (16,17).

In sequenced *V. cholerae* strains, most structural and regulatory T6SS components are encoded on a single locus on chromosome II, referred to as the large cluster (Fig. 1b). Additional components, including Hcp proteins, are encoded on two auxiliary clusters – Auxiliary clusters 1 and 2 (Aux 1, 2 respectively). Each of the three clusters also encodes a VgrG (Fig. 1b) (12,18). The VgrG encoded on the large cluster contains an additional C-terminal domain with antibacterial (lysozyme-like) activity, while the VgrG found on Aux 1 contains an anti-eukaryotic (actin-crosslinking) C-terminus domain in some strains (8,19). Terminal genes of canonical T6SS auxiliary clusters encode a secreted effector and a cognate immunity protein. Loss of immunity proteins makes cells susceptible to T6SS attacks from neighboring siblings (20,21). Both auxiliary clusters also encode TAPs (T6SS Adaptor Proteins) that are thought to be critical in loading specific effectors onto the T6SS apparatus and have been used as genomic markers to identify novel T6SS effectors (15,22). An additional cluster discovered later, Aux 3, lacks *hcp, vgrG*, and *tap* open reading frames, but contains genes coding for an effector (*tseH*) and an immunity protein (*tsiH*) (23). The Aux 3 cluster also contains a *paar* gene whose product allows the effector to be secreted by the VgrG of another cluster for delivery (23).

Regulation of T6SS genes in *V. cholerae* varies. Clinical strains, such as C6706 and A1552, show little T6SS activity in rich growth medium (24–26). Expression of genes encoded on the large T6SS cluster is up-regulated by the QstR protein, which integrates signals from three other regulators: CytR (responding to nucleoside starvation), HapR (responding to quorum sensing signals) and TfoX (responding to chitin oligomers) (27–30). By contrast, the majority of *V. cholerae* that have no history of human pathogenicity (referred to as environmental strains) express the T6SS constitutively in rich growth medium and can kill other bacterial cells in a contact-dependent manner (25). The regulation (if any) of the T6SS genes in those strains is currently not understood.

## Results

### Genome sequencing and assessment of diversity across isolates

Average nucleotide identity (ANI) was used to assess the genetic variation for environmental and clinical strains of *V. cholerae* from this study and for publicly available *V. cholerae* genomes from NCBI (31). Strain and assembly information are summarized in Supplementary Table 1. ANI revealed six clusters of *V. cholerae* strains, with clinical strains clustering together and environmental strains forming several distinct clusters (Fig. 2). SIO (BH2680), the out-group, had ANI values close to 0.96 and is at the edge of the *V. cholerae* species boundary.

**Figure 2.**
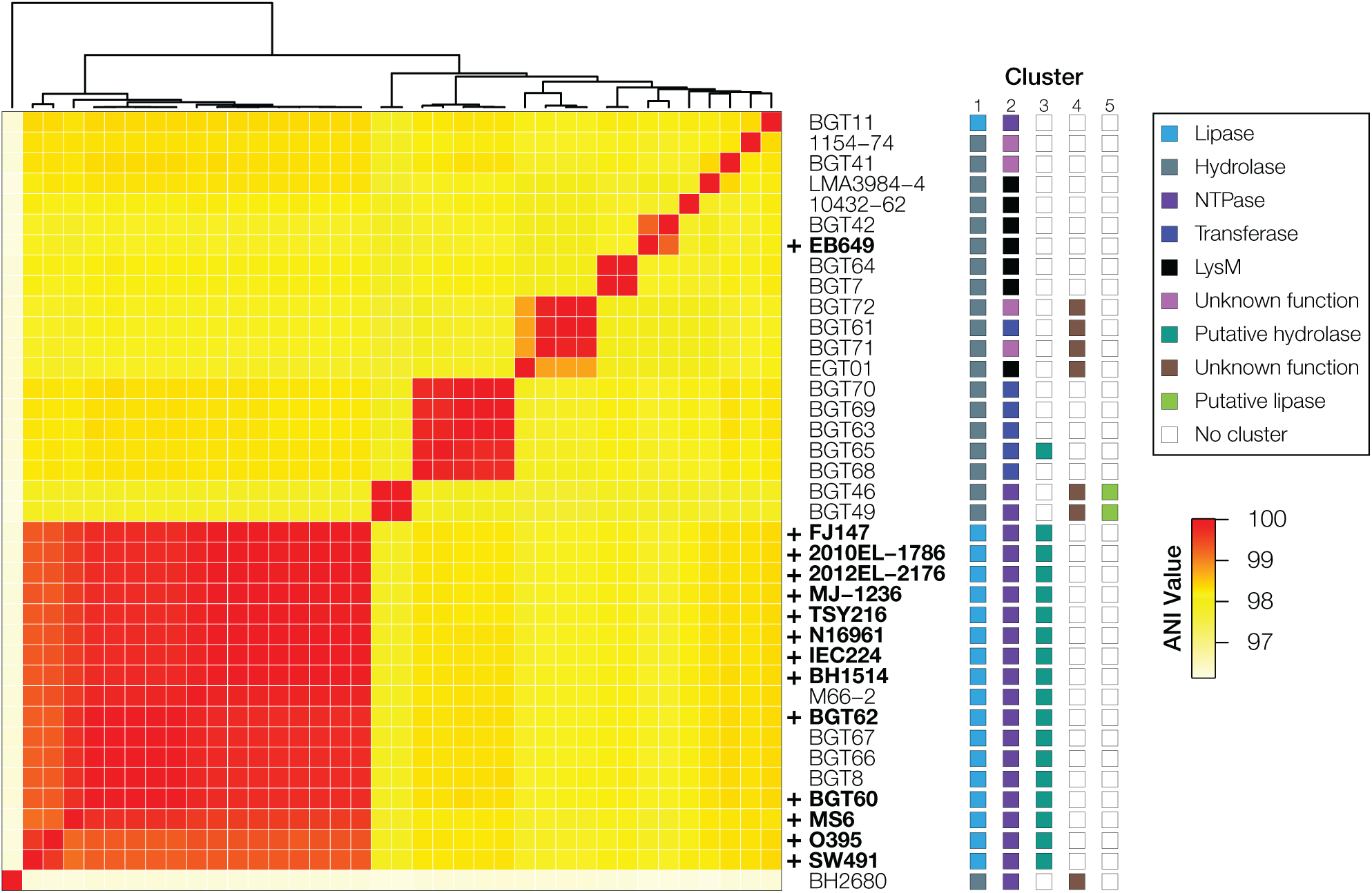
Groups of similar strains and a wide diversity of T6SS effectors are found in sequenced clinical and environmental *V. cholerae* strains. The ANI matrix includes 25 strains from this study and 14 high quality publicly available reference *V. cholerae* genomes from NCBI. ANI one-way and reciprocal best hit were used to determine protein identity between sequences. Strains which cluster together share similar phenotypes and Type VI Secretion effector-immunity proteins. A “+” sign in front of the strain name depicts the respective strain possesses the gene encoding the cholera toxin.

### T6SS module typing and annotation

Canonical *V. cholerae* T6SS loci have conserved synteny, which was used to localize searches around *vgrG* sequences to reduce the required number of BLAST searches. Initial annotation using BLAST against previously reported effector sequences was partially successful (23,32). Large, Aux 1 and Aux 2 *vgrG* alleles were successfully annotated in most strains, with occasional misannotation of *vgrG-1* alleles as *vgrG-2* and vice versa. Using this approach, we confirmed the presence of all three canonical T6SS loci (the large, Aux 1 and 2 clusters) in all sequenced isolates. By contrast, Aux 3 is found in 75% of the clinical reference strains utilized but was only observed in 30% of the isolates sequenced in this study (Fig. 2).

The conserved gene order was then used to aid in effector assignment and to identify several putative novel effectors for Aux 1 and 2. All effectors were typed and placed into classes based on conserved structural and/or functional domains (Fig. 2). T6SS effector proteins in Aux 1 were classified as lipases or hydrolases (with a DUF2235 domain). Most Aux 2 effectors were assigned as NTPases, transferases and “LysM-like” proteins. Several Aux 2 effectors (found in strains 1154-74, BGT67, BGT71 and BGT72) contain no conserved domains for typing and are dissimilar from other reported effectors and were denoted as having an “unknown function” (Fig. 2).

### Hidden Markov Models for effector prediction and annotation of new T6SS loci

To investigate whether sequenced *V. cholerae* strains contain additional, non-canonical T6SS loci, Hidden Markov Models (HMMs) were built for degenerate *hcp, vgrG* and DUF2235 hydrolase domains. Using a degenerate *hcp* HMM, an additional *hcp*-like allele was identified in six environmental strains: BGT46, BGT49, BGT61, BGT71, BGT72 and EGT01. The degenerate *vgrG* HMM identified an additional pseudo-*vgrG* in the same six strains, in-frame and directly downstream of the *hcp*-like CDS. Furthermore, the gene directly downstream of the pseudo-*vgrG* contains a DUF4123 domain found in *tap* genes. Predicted effector, immunity and *paar* genes were also observed downstream of the *tap* gene. A similar cluster previously identified in other *V. cholerae* isolates was annotated in this study as Auxiliary cluster 4 (Aux 4) (33). Aux 4 is distinct in structure, content and genomic localization from Aux 3 and is present in strains containing both Aux 1 and Aux 2 clusters.

### T6SS Predictor: web tool for prediction of V. cholerae specific T6SS protein

We also developed a tool for rapid prediction and annotation of putative T6SS loci and proteins. T6SS Predictor utilizes the profile HMMs developed for Hcp, VgrG, TAP and proteins from each effector class to annotate cluster components individually. Genomic localization and low-stringency BLAST searches using consensus sequences for each cluster/effector combination are used to assign predicted proteins to a particular cluster. Effectors are annotated using a combination of profile HMM-typing and BLAST to the custom conserved domains database used in this study. The large cluster is not annotated by T6SS Predictor. In our testing, using strains sequenced in this study, strains from Unterweger et al., and the other reference strains used in this study (Fig. 2), T6SS Predictor reliably predicts and annotates Aux 1, 2 and 3 in clinical and environmental strains and predicts Aux 4 and 5 VgrG proteins and effectors in environmental strains (32). T6SS Predictor attempts to return visualizations of each locus annotated, however contig breaks sometimes prevent the correct ordering of proteins. As a result, an annotated FASTA file containing all the predicted, putative T6SS components is provided as well.

### Aux 5 clusters have an atypical genomic organization

A profile HMM model constructed for Aux 1 DUF2235 effectors (hydrolases) identified a new putative T6SS loci in two related strains (BGT46, 49, Fig. 2). This cluster is annotated as Auxiliary cluster 5 (Aux 5) and is distinct in content and genomic organization from Aux 1, 2, 3, and 4. Aux 5 is present in *V. cholerae* strains that encode the Aux 1, 2, and 4 clusters (Fig. 2, colored boxes). The genomic organization of Aux 5 clusters is different than that of canonical T6SS auxiliary clusters in *V. cholerae* (Fig. 3a). Specifically, no open reading frames are found immediately downstream of predicted Aux 5 effectors. Instead, two genes containing DUF3304 domains found in other T6SS immunity proteins are present upstream of each effector gene.

**Figure 3.**
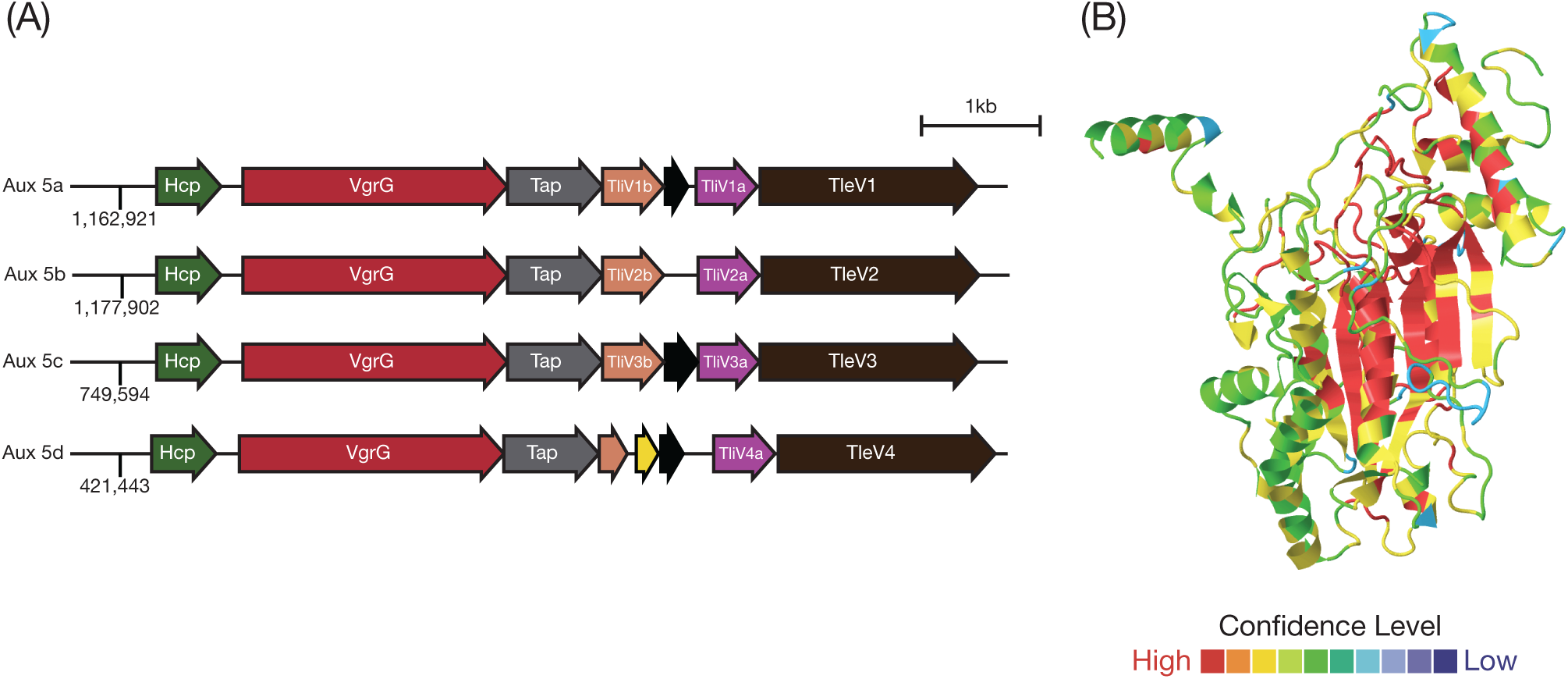
The Aux5 cluster contains an atypical gene organization and the putative effector encoded within it is predicted to be a lipase based on its structure. A) The novel Aux 5 cluster sequences from the four distinct genomic loci were aligned. The atypical Aux cluster organization is observed in all four Aux 5 clusters. Numbers at the beginning of the clusters represent the genomic position on the BGT49 chromosome. B) The structure of TleV1 was predicted using the Phyre2 webserver. The *Pseudomonas aeruginosa* Tle1 crystal structure is predicted with high confidence to be homologous to the Aux 5 putative effector. The color scheme depicts the alignment confidence of the Phyre2 model to the Tle1 crystal structure. The image was obtained using JSmol.

PacBio sequencing of strain BGT49 identified an Aux 5 cluster at four distinct genomic locations (Fig. 3a). All four Aux 5 loci (Aux 5a, b, c, d) have the same gene organization and share more than 93% nucleotide homology (Fig. 3a).

### TleV1 is toxic to E. coli cells and can be used in inter-species T6SS-mediated competition

Each predicted effector encoded within the four Aux 5 clusters contains a DUF2235 hydrolase domain, found in other T6SS-associated effectors from *Pseudomonas aeruginosa, Escherichia coli* and *Burkholderia thailandesis* (34). Phyre2 predicts with high confidence the putative effector found on cluster Aux 5a is a homologue of T6SS effector Tle1 from *P. aeruginosa*, despite sharing only 19% primary sequence identity (Supplementary Fig. 1, Fig. 3b) (35). These results reveal that the effectors belong to the larger family of Tle1 lipases that can target phospholipids and destabilize membranes. We named the putative effectors found the Aux 5 cluster TleV 1-4 (**<u>T</u>**ype VI **l**ipase **<u>e</u>**ffector **<u>V</u>**ibrio 1-4) (Fig 3a).

To experimentally validate the activity of the Aux 5a cluster, the toxicity of TleV1 was first assessed. The wild-type *tleV1* gene was expressed in *Escherichia coli* cells under the control of the arabinose-inducible pBAD promoter. Based on the predicted structure and previous similar studies showing that Tle1 lipases have activity when delivered to the periplasm, TleV1 was also expressed in *E. coli* cells with an N-terminal periplasmic Tat (twin-arginine translocation pathway) signal (34,36). When its expression was induced by arabinose, TleV1 was most cytotoxic when delivered to the periplasm, but also had moderate toxicity in the cytoplasm (Fig. 4a).

**Figure 4.**
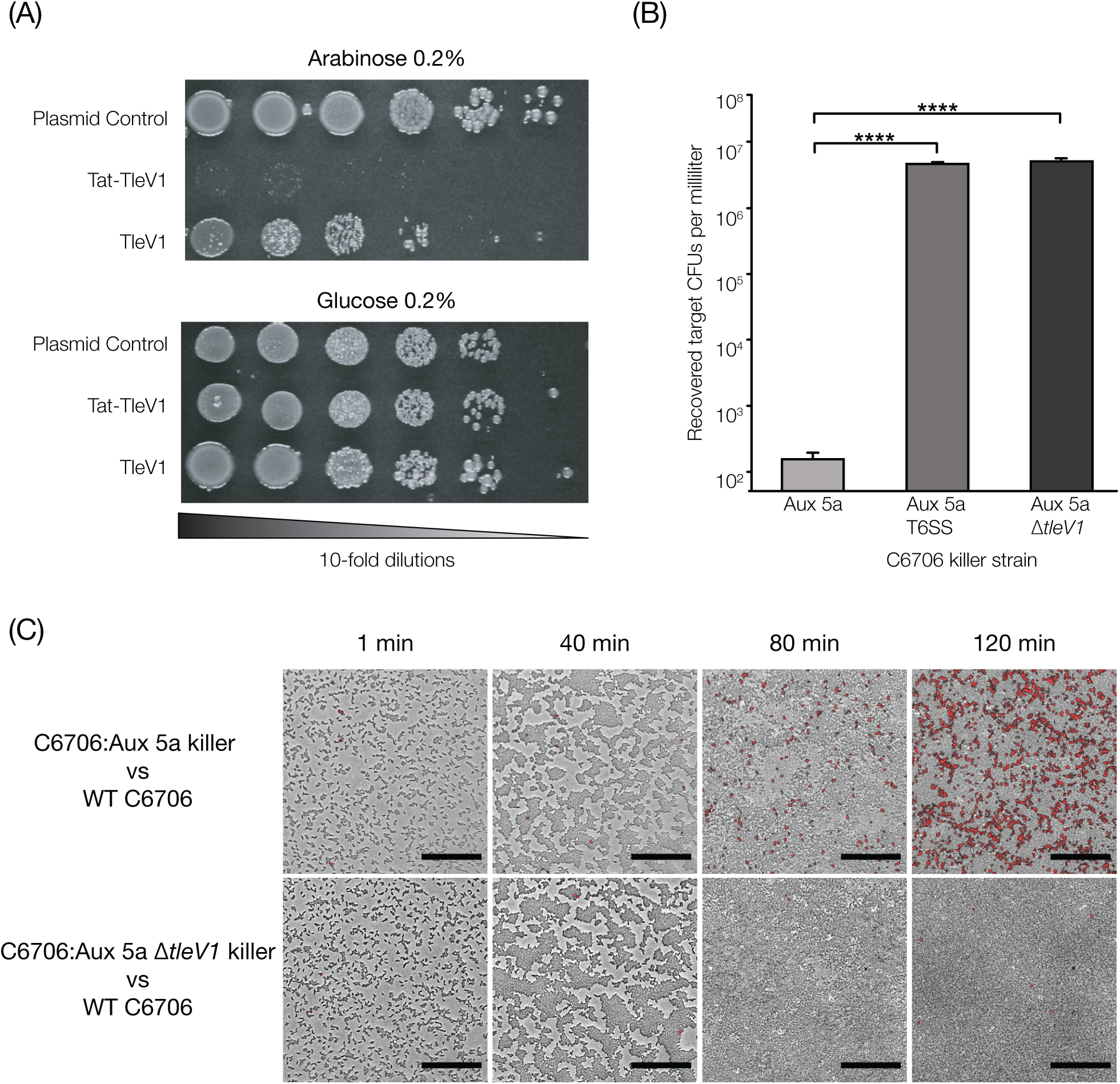
TleV1 is toxic to both *E. coli* and *V. cholerae* cells. A) Wild-type and periplasmic Tat-tagged *tleV1* genes were expressed in *E. coli* cells under the control of the pBAD promoter. Cells carrying the effector were then spotted on glucose 0.2% or arabinose 0.2% plates (and antibiotic to maintain the plasmid). B) The clinical wild-type C6706 *V. cholerae* strain was competed with C6706 with an integrated Aux5a cluster (C6706:Aux 5a) at its *lacZ* gene locus. A T6SS^−^ C6706:Aux 5a mutant and a C6706:Aux 5a Δ*tleV1* mutant were also competed against the WT C6706 target. A One-way ANOVA with post-hoc Tukey HSD Test was performed, **** p < 0.0001. C) Competitions between WT C6706 vs. C6706:Aux 5a and WT C6706 vs. C6706:Aux 5a Δ*tleV1* were visualized using propidium iodide (staining red cells with a compromised membrane) as an indicator for cellular lysis. Black scale bars represent 40 µM.

To determine whether TleV1 can be loaded onto the T6SS and be delivered to target cells, the entire Aux 5a cluster was integrated by allelic exchange methods into the *lacZ* gene locus of a clinical *V. cholerae* C6706 strain that expresses the T6SS constitutively, because the gene encoding QstR is under the control of the Ptac constitutive promoter. A competition killing assay was then performed using the *V. cholerae* C6706 strain with the integrated Aux 5a cluster (C6706:Aux 5a) as the killer strain and wild-type C6706 as the target strain. C6706:Aux 5a outcompeted wild-type C6706 and reduced the number of surviving wild-type C6706 by almost 5 orders of magnitude (Fig. 4b). A C6706:Aux 5a strain with a deletion in the essential T6SS membrane complex *vasK* gene was unable to outcompete wild-type C6706, showing that Aux 5a-mediated killing was T6SS-dependent (37). Furthermore, when *tleV1* was deleted from C6706:Aux 5a, the strain was also unable to outcompete wild-type C6706.

The presence of four Aux 5 clusters at different genomic locations suggests they may be horizontally transferred. To test this hypothesis, a Kanamycin resistance cassette was introduced immediately downstream of the Aux5a gene cluster in BGT49 using chitin-induced natural transformation. Genetic manipulation of the BGT49 is difficult because the strain was refractory to plasmid uptake by standard methods like mating or electroporation. Kanamycin-marked BGT49 genomic DNA was then used in a second natural transformation event to integrate the Aux 5 cluster into the genome of C6706. The C6706 strain containing the Aux 5 cluster was then able to successfully kill wild-type C6706 strain in a T6SS-dependent manner (data not shown). However, we observed that during transformation more than one Aux 5 cluster was transferred to the C6706 strain.

To determine whether TleV1 is toxic to cells in a manner consistent with a lipase, we examined killing induced by TleV1 using confocal microscopy (Nikon A1plus). Propidium iodide, which stains the DNA of dead cells with a compromised membrane, was used to observe cell lysis. A large number of dead cells were detected when C6706:Aux 5a killer cells were mixed with target wild-type C6706 cells (Fig. 4c). A few dead cells were observed at low cell density with little cell contact, but substantial killing occurred after two hours, when cells became densely packed. By contrast, in competitions where killer C6706:Aux 5a cells had a Δ*tleV1* deletion, only an occasional dead cell was detected throughout the time period. This result suggests TleV1 acts as a bactericidal effector when delivered into target cells.

### TliV1a can neutralize the toxic effects of TleV1

Unlike other T6SS auxiliary clusters in *V. cholerae*, where a single immunity gene is usually found downstream of an effector gene, two alleles coding for predicted immunity proteins were found upstream of each effector in all four Aux 5 clusters. For Aux 5a, we named the two genes upstream of the *tleV1* effector as *tliV1a* and *tliV1b* (**<u>T</u>**ype VI **l**ipase **i**mmunity **<u>V</u>**ibrio 1a and 1b) (Fig. 3a). To test whether the immunity gene encoded directly upstream of *tleV1, tliV1a*, can prevent self-intoxication of *E. coli* cells expressing TleV1, wild-type TliV1a or periplasmically-directed Tat-TliV1a were expressed from a second plasmid in the same cells under the control of the Ptac promoter. Survival of *E. coli* cells expressing both Tat-TleV1 and TliV1a, or both Tat-TleV1 and Tat-TliV1a, was comparable to survival of cells containing control plasmids, indicating that co-expression of the immunity gene can neutralize the toxicity of TleV1 (Fig. 5a), as shown for other effector-immunity pairs (21,32).

**Figure 5.**
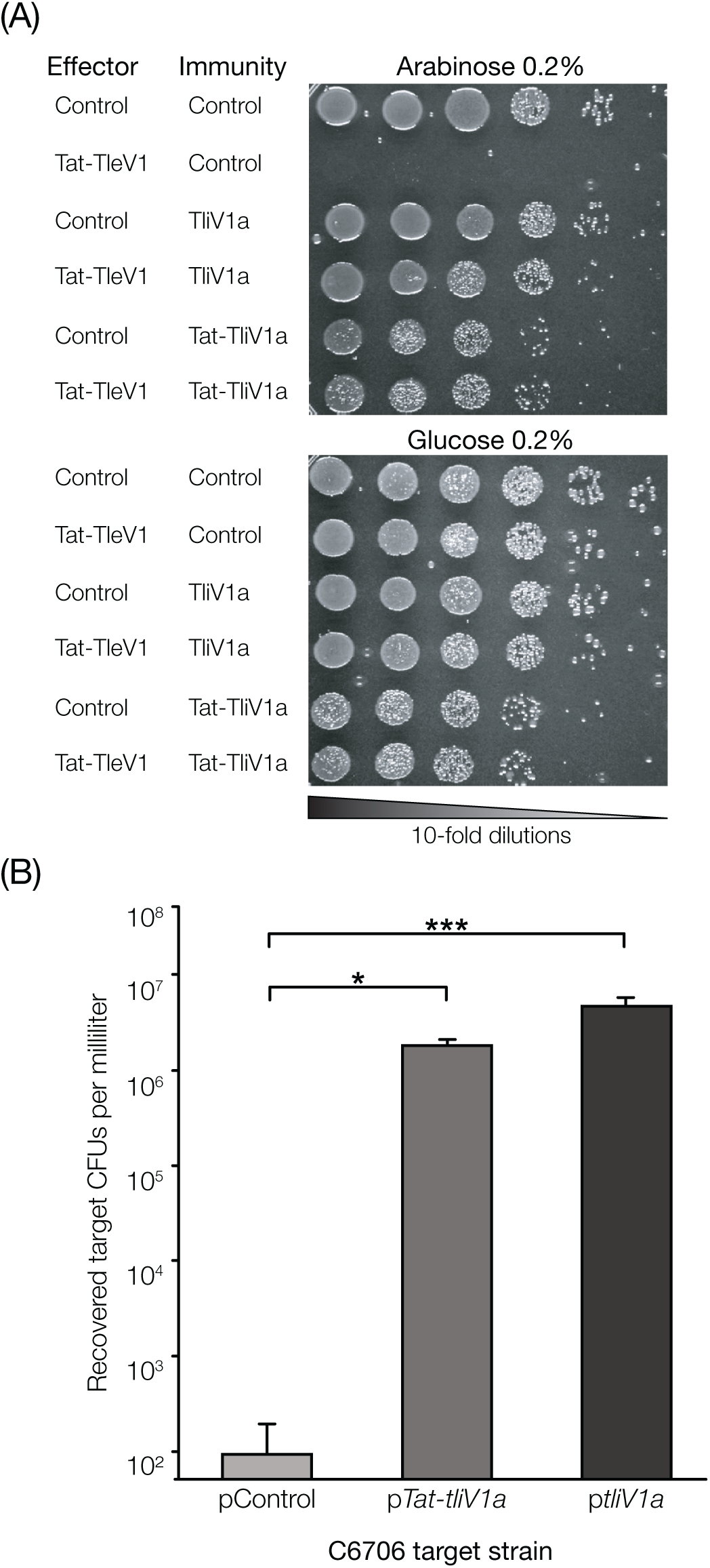
TliV1a acts as an immunity protein and neutralizes the toxic effects of TleV1. A) *E. coli* cells expressing both *Tat-tleV1* and either *tliV1a* or *Tat-tliV1a* were grown on glucose 0.2% and arabinose 0.2% (and respective antibiotics to maintain both plasmids). B) Survival of target C6706 cells carrying a plasmid control, p*tliV1a* or p*tat-tliV1a* after being competed with killer C6706:Aux 5a. A One-way ANOVA with post-hoc Tukey HSD Test was performed, *** p < 0.001, * p < 0.05.

To confirm that TliV1a can behave as an immunity protein, both TliV1a and Tat-TliV1a were expressed in wild-type C6706 *V. cholerae*. C6706 expressing either TliV1a or Tat-TliV1a was then competed as a target with killer C6706:Aux 5a. Expression of either TliV1a or Tat-TliV1a significantly rescued survival of C6706 cells as compared to C6706 cells expressing a control plasmid (Fig. 5b). These results indicate TliV1a can act as an immunity protein and prevent cellular intoxication caused by TleV1.

## Discussion

### Whole genome comparison and diversity assessment

Average Nucleotide Identity (ANI) has replaced DNA-DNA Hybridization as the species typing tool in the genomic era (38). BLAST based ANI (ANIb) has a strict species cutoff, with ANIb values < 0.96 indicating different species (38–40). As expected, clinical strains of *V. cholerae* clustered together likely due to their clonal nature (41–43). ANIb values greater than 99% are often used for subspecies or strain delineation, further supporting the clonal origins of clinical samples (44). Dot plots between strains in different ANI clusters show few, small (<20kb) rearrangements and many, small unique genomic regions (data not shown), consistent with frequent horizontal gene transfer, perhaps due to natural competence in *V. cholerae* (25,45,46).

Although *V. cholerae* strains BGT61, 71 and 72 are genetically similar and were collected in the same year (1978), they were isolated from locations more than 5000 miles apart (Supplemental Table 1). The results suggest *V. cholerae* may be widely distributed by environmental and human factors into diverse environmental reservoirs. EGT01 is genetically similar to BGT61, 71 and 72 but was collected 33 years later (2011) from grey water (water from non-sewage, home water sources) in Haiti following the 2010 cholera outbreak yet shares many of the same genomic features. EGT01 also encodes two bacterial CRISPR systems absent in the other strains, including one upstream of a T6SS cluster (31).

### Comparison with other T6SS annotation methods

Prior studies noted the difficulty involved with accurate identification and classification of diverse T6SS proteins. Unterweger et al. used a common approach, “uclust-then-BLAST”, in which predicted proteins are clustered (typically by 95% identity) followed by bi-directional best-hit BLAST searches (32). This technique is well suited for gene finding and annotation of well-characterized, conserved sequences. BLAST based approaches, as used in this study and by Unterweger et al., are also able to accurately annotate sequences with high conservation, allowing for rapid identification of canonical T6SS proteins (32). However, because of BLAST’s reliance on direct sequence comparisons and relatively high stringency in matching criteria, this approach is not well suited for exploratory annotation, especially in cases where large sequence divergence is expected. Less stringent BLAST searches can produce tens of off-target hits, such as the many transmembrane proteins that partially match VgrGs, which require significant manual curation. Manual curation is further complicated due to contig breaks, which can make unambiguous assignments of putative loci more difficult.

An existing annotation tool, SecReT6, adopts a similar clustering and BLAST approach with the addition of profile HMMs for rapid sussing out prior to BLAST (47). SecReT6’s T6SS effector database only contains T6SS alleles from clinical strains but as shown by this study and Unterweger et al., clinical strains typically contain the same effectors at Aux 1 and 2 (32). Thus, using clinical strains as the basis for effector typing under-represents the known sequence diversity of effector proteins and restricts annotations by SecReT6 to a limited set of V. *cholerae* effectors. The database contains 76 secreted effector proteins, covering the Large cluster VgrG, Lipase class Aux 1 proteins and NPPase/Transferase class Aux 2 proteins. SecReT6 is unable to identify T6SS loci in the environmental strains in this study without Lipase or NPPase Aux 1 and 2 effectors, respectively, and does not detect Aux 4 or 5 effectors. Additionally, such tools are unable to provide annotations of divergent structural proteins, such as the *hcp* and v*grG* alleles found in Aux 4 and 5, and the effectors at those loci, preventing their discovery.

The classification approach taken here differs from that used by Unterweger et al., which relied on comparing relatively large “effector modules” containing multiple variable proteins (C-terminus of VgrG, TAP, effector, and immunity), instead of comparing like to like (e.g. TAP proteins to other TAP proteins) (32). Unterweger et al. classified Aux 1 and 2 effectors into two and five categories, respectively, and the Large cluster VgrG into 7 categories. Our analysis suggests there are two Aux 1 and four Aux 2 categories based on the predicted effector activity.

### Discovery, characterization and validation of novel T6SS gene loci

All clinical *V. cholerae* isolates sequenced to date contain the same three or four T6SS genomic loci (a large cluster and two or three auxiliary clusters) and the variability of effector sequences within clinical *V. cholerae* strains is limited. By contrast, the sequenced environmental strains described here and by Bernardy et al. contain a wider diversity of effector sequences in both auxiliary clusters (25). HMM models based on degenerate *hcp* and *vgrG* genes revealed novel T6SS gene loci in environmental strains.

The Aux 4 cluster contains a canonical T6SS auxiliary cluster gene order and encodes a predicted effector. A TMHMM prediction found no transmembrane helices and predicted the effector to be non-cytoplasmic. SWISS MODEL and Phyre2 do not predict any significant homology to known structures for Tse4 but I-TASSER suggests the effector could adopt a similar fold to pilin proteins found in *Streptococcus* species (35,48,49). The activity of Aux 4 is beyond the scope of this study but is currently being investigated.

The novel Aux 5 T6SS cluster present in two sequenced *V. cholerae* strains (BGT46 and BGT49) was identified using a DUF2235 HMM. The cluster is also found in *V. cholerae* strains BK1071 (accession number GCA_900185995.1) and S12 (accession number GCA_001735565.1), which were not analyzed in this study. Short-read Illumina-based genome assembly of strain BGT49 was insufficient to resolve the gene order of the Aux 5 cluster. Subsequent sequencing of BGT49 using long-read PacBio technology confirmed the presence of *hcp, vgrG* and *tap* open reading frames and confirmed this locus is not an assembly artefact. In BGT49, sequences with high homology related to the Aux 5 cluster are found at four distinct genomic locations. The four Aux 5 genetic loci each encode a predicted effector carrying a DUF2235 hydrolase domain, found in other lipases. The genetic organization of the novel cluster is different from other *V. cholerae* T6SS clusters. The Aux 5 clusters contain two putative immunity genes containing a DUF3304 domain upstream of the putative effector. A truncated vestigial gene with limited sequence homology to *tleV* genes is also observed between the two immunity genes at all four Aux 5 loci. Phyre2 and I-TASSER predict TleV1 is most similar to Tle1 from *P. aeruginosa*, suggesting TleV1 belongs to the Tle1 family of T6SS lipases (34,35,48). TleV1 and the other three TleV alleles lack a GXSXG conserved catalytic motif associated with Tle1 lipases but contain a GXDLG motif (34).

Expression of TleV1 in the cytoplasm induced moderate toxicity in *E. coli* cells, but TleV1 was highly toxic when expressed in the *E. coli* periplasm, consistent with its designation as a Tle1-like lipase. This effect could be observed because TleV1 has catalytic activity when present in both the cytoplasm and the periplasm. Alternatively, TleV1 could have a cryptic signal that exports the wild-type protein to the periplasm even in the absence of an exogenous signal, as proposed for other T6SS effectors (50). Expression of the predicted immunity gene upstream of TleV1 was able to neutralize the toxicity of the effector in both *E. coli* and *V. cholerae* cells. Both the cytoplasmic and the periplasmic versions of TliV1a were able to rescue survival of both *E. coli* and *V. cholerae* cells. It is possible that the immunity factor is not transported to the periplasm when Tat-tagged, or may also act in the periplasm, although its transport mechanism into that compartment remains unknown.

Thomas et al. have previously shown experimentally that different effectors within *V. cholerae* auxiliary clusters can be swapped among strains (51). Kirchberger et al. have also proposed that effector modules and *tap* genes can be swapped and acquired (52). However, to our knowledge, this study is the first to experimentally show that an additional non-native T6SS auxiliary cluster can be acquired and used by a *V. cholerae* strain to kill kin cells lacking the immunity protein.

## Methods

### Vibrio cholerae culture conditions, DNA extraction and sequencing

Strains were grown overnight in LB Medium (Difco) at 37°C, with shaking. Bacterial cells were pelleted by centrifugation and the supernatant discarded. Genomic DNA was isolated using ZR Fungal/Bacterial DNA MiniPrep kit (Zymo Research) and paired-end fragment libraries constructed using Nextera XT DNA Library Preparation Kit (Illumina) with a fragment length of 300bp. For PacBio sequencing, DNA from cultures of *V. cholerae* strain BGT49 was extracted using the PacBio Phenol-Chloroform recommended protocol and cleaned using AMPure XP beads (Beckman Coulter). Purified DNA was sent to the University of Washington PacBio Sequencing Services. Raw reads were trimmed and assembled using Canu, which is designed for long read sequencing (53). Resulting contigs were then scaffolded using SSPACE-LongRead, and read correction was performed with short-read data from Illumina sequencing using Pilon (54,55).

### Genome sequence analysis

Strain and assembly information are summarized in Table 1.

### Publicly available genome sequences

Completed and publicly available *Vibrio cholerae* genome sequences were downloaded from National Center for Biotechnology Information’s (NCBI) RefSeq sequence collection and additional incomplete genomes and sequence reads archives were retrieved from NCBI’s GenBank collection and Pathosystems Resource Integration Center (PATRIC) (56–58). GenBank and RefSeq accessions are listed in Supp. Table 1.

### Whole genome comparisons

RefSeq genomes and genomes from this study were subjected to an all-by-all nucleotide comparison using one-way, reciprocal best hits BLAST to calculate percent identity between 1024bp blocks generated from each genome sequence (59). The average nucleotide identity by BLAST (ANIb) was computed for each one-way, pairwise comparison and the lower ANI value for each given pair was retained (38–40). A 30×30 symmetric matrix of ANIb values was constructed and hierarchically clustered by complete linkage and heatmap generated in R using the ggplot2 package (60,61).

### Computational characterization of T6SS

Initial identification and annotation of Large and Auxiliary T6SS clusters was by BLAST against a database constructed using sequences reported previously by Unterweger et al. and Altindis et al. (23,32). VrgG-3 and VrgG-1 and 2 alleles served as markers for putative Large and Auxiliary clusters, respectively. BLAST hits to effector proteins were considered true positives if within 3 CDS of a predicted VgrG and in the same orientation. When VgrG proteins were identified with no neighbouring effector annotation, genes at the +2 and +3 CDS with respect to VgrG protein, were marked for manual validation. All BLAST identified loci were manually validated, with new *vgrG* and effector alleles incorporated into the BLAST database. This iterative method was applied until no additional clusters were found.

Putative effector functional annotations were assigned based on conserved functional domains. Reverse, position specific BLAST (rpsBLAST) against the Protein Families (Pfam), Cluster of Orthologous Groups (COG), and Conserved Domain Database (CDD) databases was used to identify characteristic domains (62–64).

### Hidden Markov Models for effector prediction and annotation of new T6SS loci

HMMs were trained on manually curated alignments of *hcp, vgrG, tap*, effector and immunity gene sequences for each cluster type using sequences from Unterweger et al. and the strains sequenced in this study. Two additional models were created and trained for both *hcp* and *vgrG*, using sequences from other bacterial genera and *in silico* mutated sequences, respectively. HMMs were validated by reannotating the genomes from the study.

### T6SS predictor

T6SS Predictor is a Shiny application built in R using custom Perl scripts to predict and annotate putative loci (60,65). T6SS Predictor takes as input either a protein FASTA file or genomic DNA FASTA file, with the option to provide a GFF annotation file instead of relying on *de novo* CDS prediction. Predictions are generated in 2-5 minutes and the resulting output includes an annotated locus map of any identified loci and a FASTA file with putative T6SS proteins. T6SS Predictor is available from this project’s homepage: https://vibriocholera.com. T6SS predictor is hosted on fault-tolerant hardware located in the US and France and served using HTTPS best practices.

### Bacterial strains

*V. cholerae* C6706 El Tor biotype O1 strain *qstR*\* constitutively expresses the T6SS apparatus genes, and C6706 *qstR*\* Δ*vasK* is deficient for T6SS apparatus function. Both strains were used for integration and natural transformation experiments. *E. coli* MG1655 with deleted arabinose metabolism genes *araBAD* was used for expression of TleV1 from an arabinose-inducible promoter. Genomic DNA from the environmental *V. cholerae* strain BGT49 (V56) was used for Illumina and PacBio sequencing and for amplifying the Aux 5 cluster by PCR. Details regarding the *V. cholerae* and *E. coli* strains used are provided in Supplementary Tables 1 and 2.

### Modified V. cholerae strains

All C6706 *V. cholerae* genetically modified strains (both insertions and deletions) were engineered using published allelic exchange techniques (66).

### Recombinant DNA techniques

Primers used in PCR experiments were obtained from Eurofins Genomics. Phusion, Taq and Q5 Polymerases (Promega and New England Biolabs) and their respective buffers were used according to the manufacturer instructions. DNA restriction nucleases were used to digest plasmids (Promega and New England Biolabs). Gibson assembly mixes were used according to manufacturer instructions to construct the plasmids used in this study (New England Biolabs). All recombinant strains and constructs used in the study were tested by colony PCR and verified for accuracy by Sanger sequencing.

### E. coli toxicity experiments

*E. coli* strains expressing the *tleV1* gene under control of arabinose-inducible pBAD promoter were cultured in LB medium with 150 µg/mL of spectinomycin and 0.2% glucose overnight. Cells were then washed three times with LB and resuspended in fresh culture medium to an OD_600_ of 0.5. To assess toxicity, 10-fold serial dilutions were performed, and 3 µL aliquots of cell suspensions were then spotted on agar plates containing either spectinomycin and 0.2% glucose or containing spectinomycin and 0.2% arabinose. Agar plates were incubated statically overnight at 37°C. The same growth conditions were used for *E. coli* strains expressing both *tleV1* and *tliV1a* genes with the exception that cells were grown overnight in LB medium with 150 µg/mL of spectinomycin, 10 µg/mL of chloramphenicol and 0.2% glucose overnight and then were spotted on agar plates containing either spectinomycin, chloramphenicol and 0.2% glucose or containing spectinomycin, chloramphenicol and 0.2% arabinose.

### T6SS Killing Assays

*V. cholerae* strains (both killer and target) were incubated overnight with shaking in liquid LB at 37°C. Both strains were then washed 3 times with LB, diluted to an OD_600_ of 1 in fresh LB and then mixed together in a 10:1 (killer:target) ratio. Aliquots (50 µL) of the mixed cell suspension were spotted on filter paper with a 0.2 µm pore size that was placed on an LB plate and incubated at 37°C for 3 hours. Each filter paper was then vortexed for 30 seconds in 5 mL of LB. Resuspended cells were diluted and spread on plates containing antibiotic to select for surviving target cells. Plates were then incubated at 37°C overnight and the number of colonies was counted.

### Confocal Microscopy Experiments

*V. cholerae* strains (both killer and target) were incubated overnight with shaking in liquid LB at 37°C. Each overnight culture was back-diluted 1:100 and incubated with shaking at 37°C for approximately 6 hours. Cell suspensions were then normalized to an OD_600_ of 1 in fresh LB and mixed in a 1:1 (killer:target) ratio. An 8 µL aliquot of propidium iodide (100 µg/mL) was added to an agar pad and allowed to dry. Next, a 1 µL aliquot of the killer:target cell mixture was spotted. Cells were imaged at 37°C and 96-100% humidity for 5 hours using an Eclipse Ti-E Nikon inverted microscope. A Perfect Focus System was used with a 40x objective (Plan Fluor ELWD 40x DIC M N1) to stabilize the focus in the plane of the biofilm growth during long-term imaging. A Nikon A1plus camera was used to obtain images. Images were processed in ImageJ.

### Natural Transformation Experiments

Natural transformation experiments were performed as described by Watve et al. (67). Briefly, *V. cholerae* overnight cultures were diluted 1:100 in fresh LB medium and allowed to achieve an OD_600_ of ∼0.3. 2 ml of each culture was then added to a sterile crab shell fragment and was incubated overnight at 30°C in artificial sea water medium (17 g/liter of Instant Ocean, cat. no. SS115-10). Genomic DNA from donor bacteria containing an antibiotic resistance gene was added and the cells were incubated for 24 hours. Cells were then spread on plates containing antibiotic to select for transformed cells.

## Conclusion

Competition within microbial communities is an important aspect of the life cycle of *V. cholerae* and other benign and pathogenic microbes. Twenty-six *V. cholerae* strains were sequenced and Hidden Markov Models were used to probe for novel gene clusters associated with T6SS activity in *V. cholerae* isolates. Using the bioinformatics tool we developed, a novel cluster, named auxiliary cluster 5 (Aux 5), was discovered and the effector encoded within the cluster was toxic when expressed in *E. coli* cells. The entire cluster was transferred into a different *V. cholerae* strain and conferred T6SS-dependent competitive advantage to the recipient strain. We propose that the tool we have developed is better suited than previous methods for discovering novel T6SS effectors in *V. cholerae* species and may be adapted to facilitate discovery of effectors in other bacterial species.

## Declarations

Authors have no completing financial or non-financial interests.

Data and material not provided in the text, such as DNA primers, are available upon request. T6SS Predictor is available from this project’s homepage: https://vibriocholera.com.

## Authors’ contributions

Cristian V. Crisan and Kenneth Williams – experimental design, all wet lab experiments.

Aroon T. Chande, Vishnu Raghuram and Lavanya Rishishwar – experimental design, all bioinformatics analyses.

Gabi Steinbach – confocal microscopy experiments.

Brian K. Hammer, I. King Jordan and Peter Yunker – experimental design, funding, advice and critiques.

## Acknowledgements

We would like to thank all current and past members of the Hammer Lab who have contributed to this paper with critiques and discussions, in particular: Samit Watve, Jacob Thomas, Harshini Chandrashekar, Siu Lung Ng.

## Figure Legends

**Supplementary Figure 1. TleV1 is predicted to share structural homology to the *P. aeruginosa* Tle1 T6SS lipase.**

Phyre2 modelling of TleV1 predicts its secondary structure is similar to Tle1, despite sharing only 19% amino acid sequence identity.

**Supplementary Figure 2. TleV1 is predicted to share structural homology to the *P. aeruginosa* Tle1 T6SS lipase.**

Annotation workflow. Graphical description of the annotation workflow used in this study and, in part, by T6SS predictor.

