## Supplementary figures and images for "Analysis of *Vibrio cholerae* genomes using a novel bioinformatic tool identifies new, active Type VI Secretion System gene clusters"

### Supplementary Figure 1

Supplementary Figure 1.

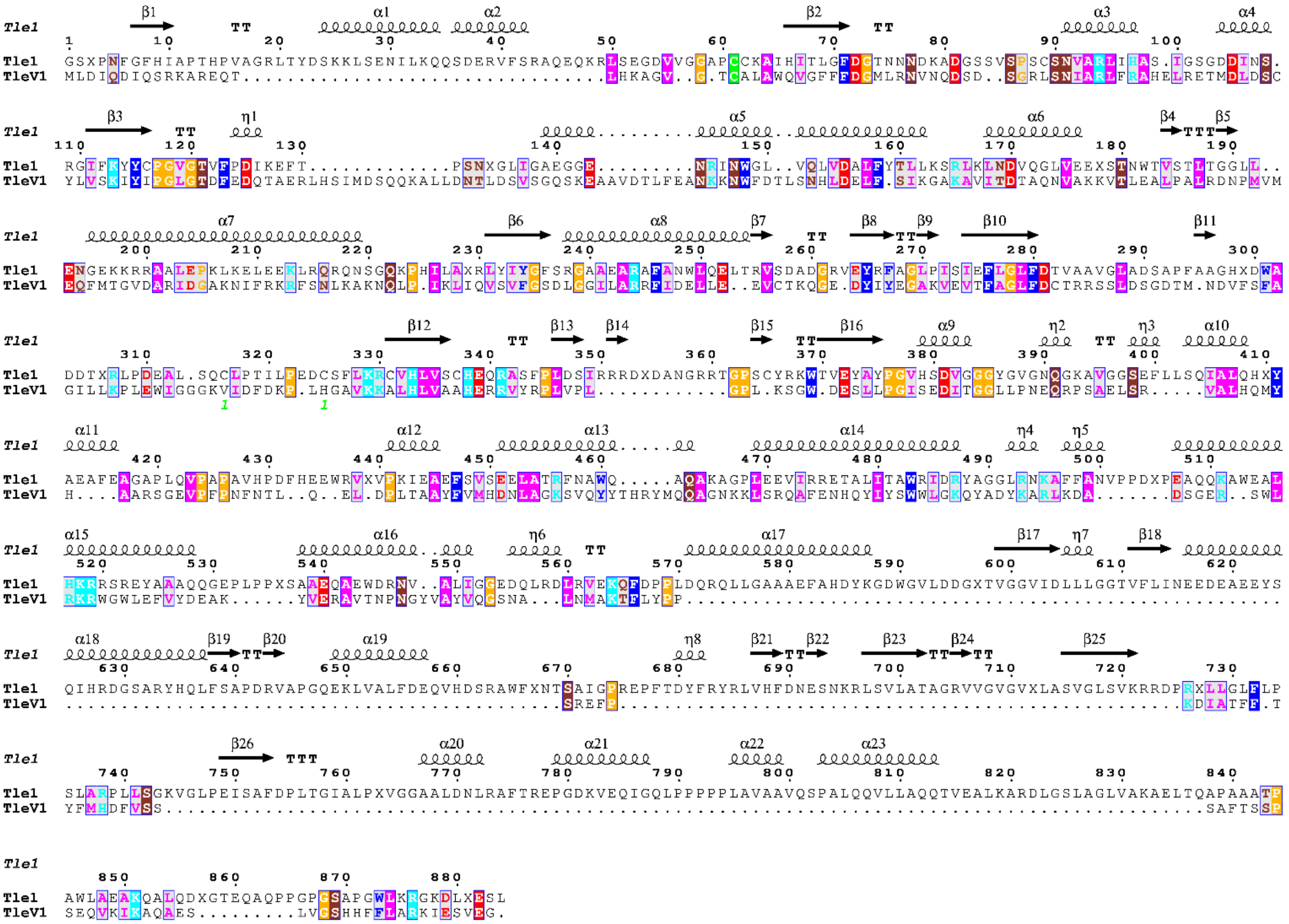

### Supplementary Figure 2

Supplementary Figure 2.

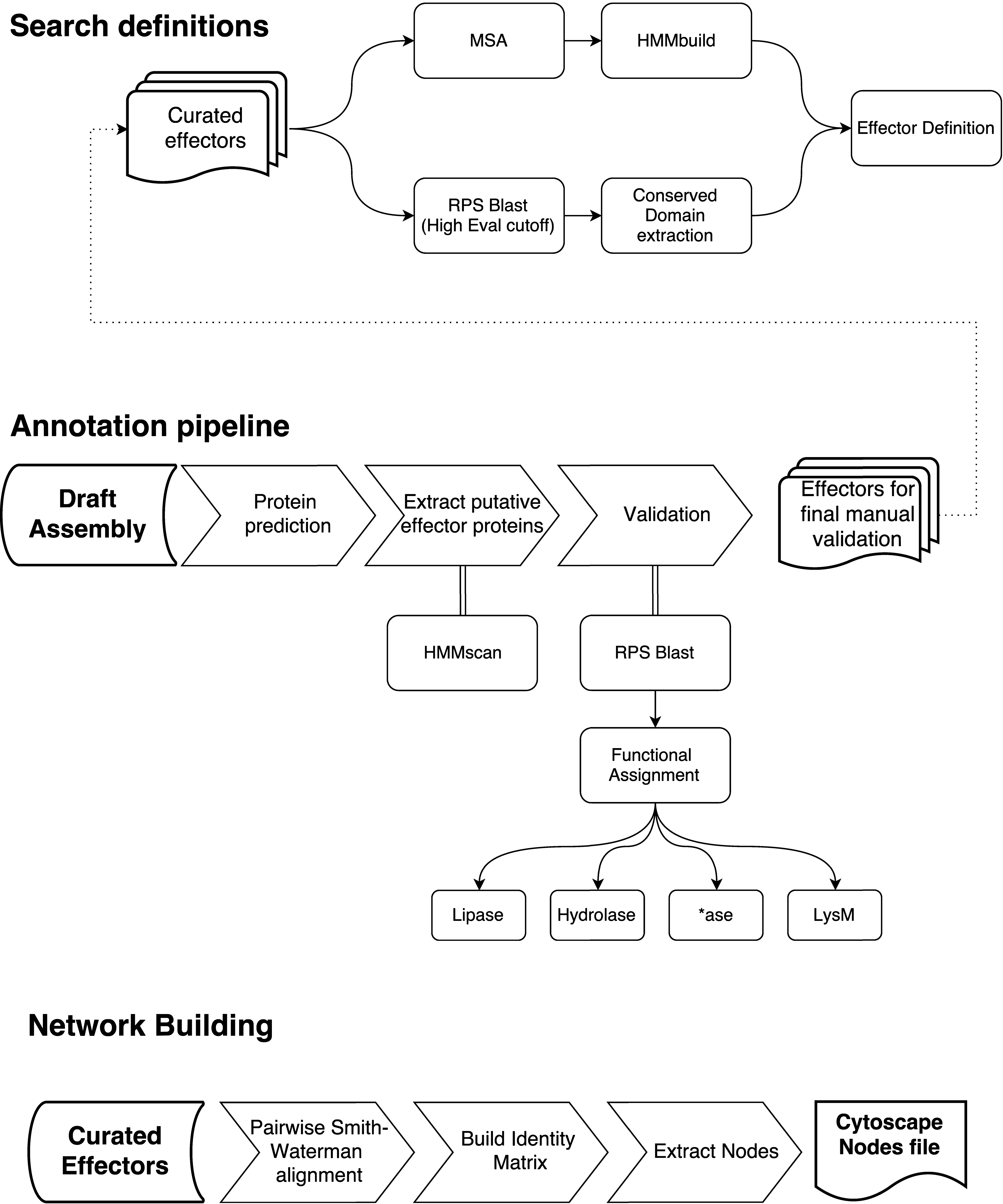
