## Supplementary Table 1 for "Analysis of *Vibrio cholerae* genomes using a novel bioinformatic tool identifies new, active Type VI Secretion System gene clusters"

| Strain | Strain number | Accession | Location | Source | CTX | Year of isolation | Type VI killing activity | Coverage | Number of scaffolds | Assembled bases | N50 | L50 | GC % | Ns/100Kbp |
|---|---|---|---|---|---|---|---|---|---|---|---|---|---|---|
| V52 | SW491 | GCA_001857545.1 | Sudan | Patient | + | 1968 | + | 2,504x | 175 | 3,957,367 | 177,630 | 7 | 47.56 | 13.52 |
| 3223-74 | BGT69 | GCA_001743085.1 | Guam | Storm drain | - | 1974 | + | 2,740x | 173 | 4,069,095 | 279,340 | 6 | 47.65 | 5.97 |
| 2523-87 | BGT68 | GCA_001857345.1 | USA (LA) | Moore swab | - | 1974 | + | 1,605x | 177 | 3,964,384 | 320,410 | 4 | 47.69 | 29.89 |
| 3225-74 | BGT70 | GCA_001857365.1 | Guam | Storm drain | - | 1974 | + | 2,451x | 203 | 4,057,051 | 387,604 | 4 | 47.79 | 4.63 |
| 3272-78 | BGT63 | GCA_001857265.1 | USA (MD) | Water | - | 1977 | + | 2,245x | 143 | 3,999,272 | 289,139 | 3 | 47.58 | 6.98 |
| 2559-78 | BGT60 | GCA_001857145.1 | USA (LA) | Crab | + | 1978 | + | 1,736x | 164 | 4,040,166 | 298,103 | 6 | 47.61 | 17.2 |
| 2631-78 | BGT61 | GCA_001857225.1 | USA (LA) | Moore swab | - | 1978 | + | 1,947x | 176 | 3,995,972 | 150,154 | 10 | 47.67 | – |
| 1074-78 | BGT71 | GCA_001857405.1 | Brazil | Sewage | - | 1978 | + | 1,644x | 175 | 4,003,881 | 127,547 | 12 | 47.68 | 21.75 |
| 2633-78 | BGT72 | GCA_001857425.1 | Brazil | Sewage | - | 1978 | + | 2,714x | 154 | 3,982,026 | 127,546 | 11 | 47.67 | 21.57 |
| 692-79 | BGT64 | GCA_001857285.1 | USA (LA) | Water | - | 1979 | + | 2,115x | 122 | 3,953,095 | 249,947 | 5 | 47.63 | 14.14 |
| 2740-80 | BGT08 | GCA_001729185.1 | USA (Gulf coast) | Water | - | 1980 | + | 2,172x | 223 | 4,040,634 | 260,656 | 6 | 47.73 | 14.35 |
| VC22 | BGT41 | GCA_001729195.1 | USA (FL) | Oyster | - | 1981 | + | 1,650x | 146 | 4,062,714 | 154,565 | 9 | 47.53 | 13.83 |
| VC48 | BGT42 | GCA_001857165.1 | USA (FL) | Oyster | - | 1981 | + | 1,612x | 176 | 3,954,170 | 114,301 | 11 | 47.57 | 14.57 |
| 2512-86 | BGT62 | GCA_001857245.1 | USA (LA) | Moore swab | + | 1986 | + | 1,380x | 212 | 4,035,883 | 298,103 | 6 | 47.69 | 18.43 |
| 2479-86 | BGT65 | GCA_001857305.1 | USA (LA) | Moore swab | - | 1986 | + | 1,766x | 17 | 4,039,476 | 320,139 | 4 | 47.69 | 4.36 |
| 2497-86 | BGT67 | GCA_001857355.1 | USA (LA) | Moore swab | - | 1987 | + | 2,314x | 159 | 4,012,321 | 298,103 | 6 | 47.57 | 13.68 |
| 857 | BGT07 | GCA_001729125.1 | Bangladesh | Water | - | 1996 | + | 1,508x | 131 | 4,020,975 | 249,952 | 5 | 47.67 | 15.17 |
| SIO | BH2680 | GCA_001857455.1 | USA (CA) | Water | - | 2000 | + | 1,504x | 173 | 4,017,200 | 200,950 | 7 | 47.19 | 2.64 |

| | | | | | | | | | | | | | |
|---|---|---|---|---|---|---|---|---|---|---|---|---|---|
| 3568-07 | EB649 | GCA_001857505.1 | Mexico | Queso fresco | + | 2007 | + | 2,366x | 172 | 4,095,080 | 106,941 | 16 | 47.37 | 2.34 |
| VC53 | BGT46 | GCA_001857155.1 | USA (AL) | Oyster | - | 2009 | + | 2,073x | 267 | 4,239,039 | 86,454 | 14 | 47.26 | 48.55 |
| VC56 | BGT49 | GCA_001857175.1 | USA (AL) | Oyster | - | 2009 | + | 1,710x | 244 | 4,227,097 | 86,314 | 15 | 47.29 | 14.22 |
| HE46 | EGT01 | GCA_001857515.1 | Haiti (Centre) | Gray water | - | 2011 | + | 2,424x | 167 | 4,039,421 | 149,375 | 11 | 47.68 | 35.04 |
| 1496-86 | BGT66 | GCA_001857325.1 | USA (LA) | Moore swab | - | 1986 | - | 1,511x | 161 | 3,982,315 | 280,655 | 6 | 47.54 | 2.26 |
| C6706 | BH1514 | GCA_001857435.1 | Peru | Patient | + | 1991 | - | 1,204x | 150 | 4,035,736 | 260,568 | 6 | 47.45 | 36.06 |
| MZO-2 | BGT11 | GCA_001729155.1 | Bangladesh | Patient | - | 2001 | - | 2,356x | 161 | 4,001,541 | 255,781 | 5 | 47.51 | 27.96 |
