## Supplementary Table 2 for "Analysis of *Vibrio cholerae* genomes using a novel bioinformatic tool identifies new, active Type VI Secretion System gene clusters"

**Bacterial Strains**

| **Strain** | **Description** | **Genotype** | **Reference** |
|---|---|---|---|
| NRD206 | *E. coli* MG1655 | *ΔlacZY ΔaraBAD* | N. R. De Lay and J. E. Cronan. J. Biol. Chem. 282.28: 20319-28, 2007 |
| BH1514 | *V. cholerae* C6706 | El Tor biotype O1 | K. H. Thelin and R. K. Taylor, Infect. Immun. 64(7): 2853-2856, 1996 |
| KW25 | *V. cholerae* C6706 | *ptac-qstR, ΔlacZ::Aux5a* | This study |
| KW26 | *V. cholerae* C6706 | *ptac-qstR, ΔvasK, ΔlacZ::Aux5a* | This study |
| CC111 | *V. cholerae* C6706 | *ptac-qstR, ΔlacZ::Aux5a, ΔtleV1* | This study |
| JT516 | *V. cholerae* C6706 | *ΔlacZ::SpecR* | This study |
| CC94 | *V. cholerae* C6706 | *ΔlacZ::SpecR, pSLS3* | This study |
| KW18 | *V. cholerae* C6706 | *ΔlacZ::SpecR, pSLS3-tliV1a* | This study |
| CC95 | *V. cholerae* C6706 | *ΔlacZ::SpecR, pSLS3-tat-tliV1a* | This study |

**Plasmids**

| **Plasmid** | **Features** | **Reference** |
|---|---|---|
| pBAD18 | *pBAD* promoter, SpecR, pSLS3 origin or transfer | L.M. Guzman, et al. J. Bacteriol. 177 4121-30, 1995 |
| pBAD18TleV1 | *tleV1a*, SpecR | This study |
| pBAD18Tat-TleV1 | *tat-tleV1a*, SpecR | This study |
| pSLS3 | CmR | K. C. Tu and B. L. Bassler. Genes& dev. 21: 221-233, 2007 |
| pSLS3TliV1a | *ptac-tliV1a*, CmR | This study |
| pSLS3Tat-TliV1a | *Ptac-tat-tliV1a*, CmR | This study |
